# Exploring the molecular composition of the multipass translocon in its native membrane environment

**DOI:** 10.1101/2023.11.28.569136

**Authors:** Max Gemmer, Marten L. Chaillet, Friedrich Förster

**Author notes:** Correspondence to David de Wiedgebouw, Universiteitsweg 99, Kamer 2.70, 3584 CG Utrecht, The Netherlands.

## Abstract

Multispanning membrane proteins are inserted into the endoplasmic reticulum membrane by the ribosome-bound multipass translocon machinery. Based on cryo-electron tomography and extensive subtomogram analysis, we reveal the composition and arrangement of multipass translocon components in their native membrane environment. The intramembrane chaperone complex PAT and the translocon associated protein (TRAP) complex associate substoichiometrically with the multipass translocon in a translation-dependent manner. While PAT is preferentially recruited to active complexes, TRAP primarily associates with inactive translocons. The subtomogram average of the TRAP-multipass translocon reveals intermolecular contacts between the luminal domains of TRAP and an unknown subunit of the BOS complex. AlphaFold modeling suggests this protein is NOMO, bridging the luminal domains of nicalin and TRAPα. Collectively, our results visualize the interplay of accessory factors associated with multipass membrane protein biogenesis under near-native conditions.

## Introduction

The ribosome-associated ER translocon complex facilitates the biogenesis of most secretory and membrane proteins ^1^. The ribosome binds to the protein-conducting channel Sec61, a heterotrimeric ER membrane-embedded complex, which facilitates signal peptide (SP) insertion into its lateral gate and co-translational translocation of the nascent chain into the ER lumen through its central pore ^2^. To meet the requirements of the broad spectrum of nascent protein clients, Sec61 associates with various accessory factors specialized in functions such as SP insertion and cleavage, N-glycosylation, protein folding and maturation, transmembrane helix insertion, and ER stress response ^1,3–6^. Sec61 assembles with different sets of accessory factors in ER translocon complexes. In all mammalian cells studied to date, the most abundant ribosome-bound ER translocon variant comprises the translocon-associated protein (TRAP) complex and the oligosaccharyltransferase A (OSTA) complex in addition to Sec61 ^7^. A less abundant variant comprises only Sec61 and TRAP. The heterotetrameric TRAP complex facilitates insertion of SPs with below-average hydrophobicity and above-average glycine-and-proline content ^8,9^, while OSTA mediates co-translational N-glycosylation and recruitment of ER luminal chaperones ^10,11^.

In contrast to secretory proteins, biogenesis of multispanning transmembrane proteins relies on different translocon variants that specialize on transmembrane helix (TMH) insertion, membrane protein topogenesis, folding and assembly ^4^. Recently, Get1, EMC3, and TMCO1 have been identified as ER resident Oxa1 superfamily members, which are core components of different insertase complexes^12^. The guided entry of tail-anchored protein (GET) complex facilitates post-translational targeting and insertion of tail-anchored membrane proteins ^13,14^, while the ER membrane complex (EMC) and the TMCO1-translocon complex facilitate co-translational TMH insertion of multispanning membrane proteins ^12,15–19^. Recent biochemical, mass-spectrometry and cryo-electron microscopy (cryo-EM) studies of affinity-tagged TMCO1 isolates revealed the composition and architecture of a ribosome-bound TMCO1-containing complex ^19–21^, herein referred to as the multipass translocon.

Besides Sec61, the multipass translocon comprises three sub-complexes: the GET– and EMC-like (GEL) complexes, the protein associated with the ER translocon (PAT), and the back-of-Sec61 (BOS) complex. The GEL complex consists of TMCO1 and obligate partner of TMCO1 insertase (OPTI), and it facilitates TMH insertion ^19–22^. The PAT complex consists of the intramembrane chaperone asterix to protect TMHs with exposed hydrophilic residues ^23^, and CCDC47, which forms contacts with the ribosome and impedes Sec61 opening ^23,24^. The BOS complex comprises TMEM147, nicalin (NCLN), and one of the three nearly identical paralogs of nodal modulator (NOMO) 1, NOMO2, or NOMO3, collectively referred to as NOMO ^25–27^. The function of the BOS complex is poorly understood. TMEM147, as well as *C. elegans* homologs of NCLN (NRA-2) and NOMO (NRA-4), affect levels and subcellular localization of different multi-spanning membrane proteins, while NOMO has been shown to play an additional role as ER sheet shaping protein ^28–30^. Together, the components of the multipass translocon form a lipid-filled cavity adjacent to Sec61, which mediates TMH insertion and folding of multi-spanning membrane proteins ^19^. While cryo-EM studies revealed the structural organization of the multipass translocon, its activity-dependent compositional variation in the native membrane remains to be explored.

Here we analyze the composition and organization of the multipass translocon in the native ER membrane using electron cryo tomograms of ER microsomes ^31^. While earlier analysis covered distinct ribosomal intermediate states, major translocon variants, and the Sec61-TRAP-OSTA translocon, we now focus on subtomogram analysis of ribosome-multipass-translocon complexes. The analysis reveals the compositional differences of the multipass-translocon dependent on translation activity and ER stress.

## Results and Discussion

We previously analyzed ∼135,000 ribosome particles from HEK293 cell-derived ER microsomes and revealed the distribution of the major ER translocon variants. We have shown that ∼14,700 ribosomes were associated with the multipass translocon variant (Fig. 1A,B). While refinement of the ribosome-multipass translocon yielded a reconstruction with a resolution of ∼8 Å in the vicinity of Sec61, its associated accessory factors remained poorly resolved (Fig. 1B).

**Fig. 1:**
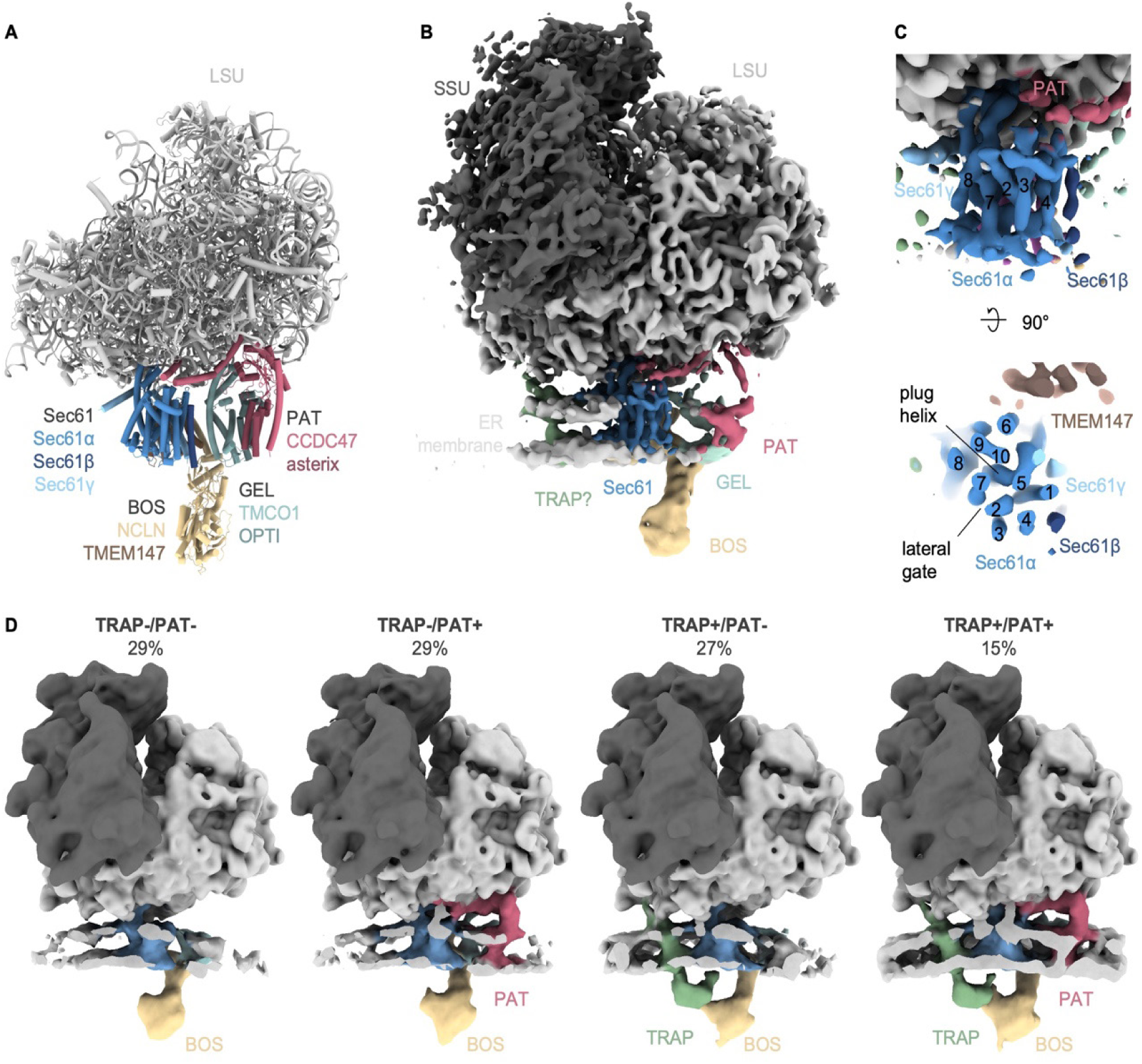
Structure and composition of the multipass translocon in the native ER membrane. (A) Structure of the ribosome-associated multipass translocon (PDB-7TUT) determined by Smalinskaite *et al.* ^20^. (B) Cryo-ET reconstruction of the heterogenous ribosome-associated multipass translocon-population (14,671 particles). Ribosome and Sec61 were filtered to 8 Å and the accessory factors and ER membrane area to 20 Å. (C) Close-up front (top panel) and top (bottom panel) view of Sec61 from the multipass translocon population from (B). Numbering of TMHs of Sec61α is indicated. (D) Reconstructions of different multipass translocon populations. Reconstructions were filtered to 20 Å resolution. The ER membrane was clipped for visual clarity.

The Sec61 lateral gate and plug helix both adopt a closed conformation that matches the structure of the isolated TMCO1 translocon ^19^ (Fig. 1C). This contrasts the conformation of the cryo-ET reconstruction of the Sec61-TRAP-OSTA-translocon, where the Sec61 complex adopts an open plug and the lateral gate accommodates a helical peptide ^31^. Thus, the fully-closed Sec61 conformation is a characteristic of the multipass translocon complex in the native membrane.

### PAT and TRAP are variable multipass translocon components

To reveal potential substoichiometric components or conformational heterogeneity, subtomograms of multipass translocon particles were subjected to two independent rounds of 3D classification focused either on the PAT complex or on luminal densities of the TRAP and BOS complexes. This classification procedure yields four classes, comprising either TRAP only (TRAP+/PAT-, 27%), PAT only (TRAP-/PAT+, 29%), both (TRAP+/PAT+, 15%), or none (TRAP-/PAT-, 29%) (Fig. 1D). Hence, TRAP and PAT are both substoichiometric subunits of the multipass translocon.

Interestingly, we observed a slight but significant reduction of PAT levels in the presence of TRAP (p=7.14•10^-12^), and, *vice versa*, a reduction of TRAP in the presence of PAT (p=8.52•10^-11^) (Fig. 2A,B). We had previously knocked out CCDC47, a subunit of the PAT complex, to identify the multipass translocon in tomograms of microsomes isolated from ΔCCDC47 cells ^31^. To further investigate the correlation of TRAP and PAT in their association with the multipass translocon, we now revisited this datatset using subtomogram classification. Remarkably, in the ΔCCDC47 microsomes, we did not observe a TRAP-less multipass translocon population (Fig. 2C,D). Since TRAP and PAT reside at opposite sides of Sec61, they do not display any notable clashes (Fig. 1) and thus do not compete for a mutual binding site. Taken together, TRAP is a stoichiometric component of the multipass translocon in the absence of PAT.

**Fig. 2:**
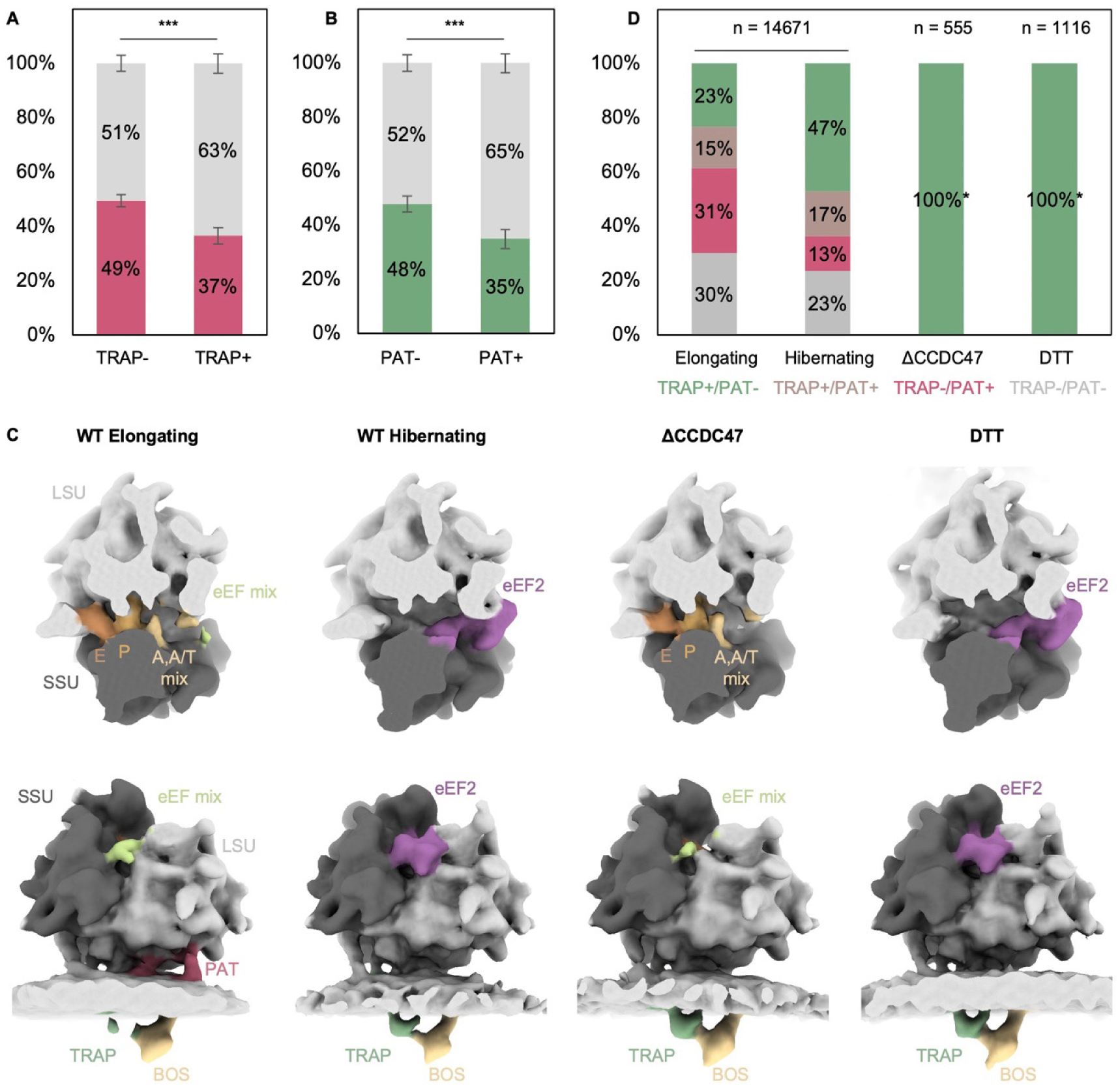
Recruitment of PAT and TRAP is translation-dependent. (A) Relative abundance of PAT (magenta) in the TRAP-lacking (TRAP-) or –containing (TRAP+) multipass population with p=7.14•10^-12^. (B) Relative abundance of TRAP (green) in the PAT-lacking (PAT) or –containing (PAT+) multipass population with p=8.52•10^-11^. (C) Reconstructions of multipass translocon-associated ribosomes obtained from HEK293F under unperturbed condition (WT) of elongating and hibernating classes, ΔCCDC47, and DTT-treated samples. Top row: top views of the ribosomal binding cleft and elongation factor binding site. The ribosome was clipped for clarity. Bottom row: corresponding front views of the ribosome-associated translocon populations. The ER membrane was again clipped for clarity. The subtomogram averages represent the entire multipass population in each class (hibernating and elongating) or sample (ΔCCDC47 and DTT). Note the varying occupancy of TRAP and PAT in different classes. (D) Distribution of TRAP and PAT in the multipass populations from (C) color-coded as indicated. (*) PAT-containing or TRAP-lacking classes were not detected in ΔCCDC47 and DTT-treated samples.

To explore the translation activity-dependent recruitment of TRAP and PAT, we analyzed the distribution of multipass translocon classes in the context of the translation state of their bound ribosomes. We previously separated active and inactive ribosomes by grouping the particles into elongating and hibernating populations using extensive classification of ribosomal intermediate states ^31^. Strikingly, in multipass translocon particles associated with inactive ribosomes, we observed a strong increase of the TRAP+/PAT-class from 23% to 47%, accompanied by a strong reduction of the TRAP-/PAT+ class from 31% to 13% (Fig. 2C,D). Hence, TRAP is preferably recruited to inactive ribosome-multipass translocon particles, while PAT is found primarily in active ribosome-multipass translocons.

To support the notions that TRAP recruitment to the multipass translocon correlates with translational inactivity and PAT with translational activity, we revisited cryo-ET data of vesicles isolated from HEK cells, which were treated with the ER stress-inducing drug dithiothreitol (DTT) ^31^. Induction of the unfolded protein response triggers a dramatic reduction of translational activity (Fig. 2C) ^32^. Remarkably, we found TRAP to be stoichiometric in the multipass translocon associated with the almost exclusively inactive ribosome (97%) ^31^, while PAT was not detected (Fig. 2C,D). We attribute this result to the preference of TRAP to associate with inactive translocons. However, the enhanced recruitment of TRAP may also be explained by ER stress-activated upregulation of all four TRAP subunits ^33^ and thus could be a direct result of ER stress. Importantly, the knockout of the PAT subunit CCDC47 does not appear to significantly affect ribosomal translation (Fig. 2C), precluding the possibility that the observed recruitment effects induced by the ΔCCDC47 are caused indirectly by reduced translational activity.

Recently, a cryo-EM single particle analysis (SPA) study revealed the architecture of the isolated, ribosome-bound Sec61-TRAP translocon ^34^. The high resolution of the ribosome-TRAP interface in this map allowed identification of a short cytosolic, C-terminal stretch of the TRAPα subunit anchored to the 5.8S ribosomal RNA (rRNA) of the large ribosomal subunit. Consistently, we spot this anchor in subtomogram averages of membrane-bound ribosome classes (Sec61-TRAP-OSTA), but not in a soluble population revealed in our previous analysis ^31^ (Fig. 3). Interestingly, we also observe a small density colocalizing with the TRAPα anchor in multipass variants regardless of the association of the core TRAP complex (Fig. 3). While the resolution is not sufficient to unambiguously identify this density, it co-localizes with the TRAPα anchor assigned in the higher resolution maps of the isolated, ribosome-bound Sec61-TRAP translocon ^34^. Consistent with our observation, a previous study proposed that the TRAPα anchor flexibly tethers the complex to the ribosome even when the transmembrane and luminal TRAP segments detach from the ribosome-translocon complex ^34^.

**Fig. 3:**
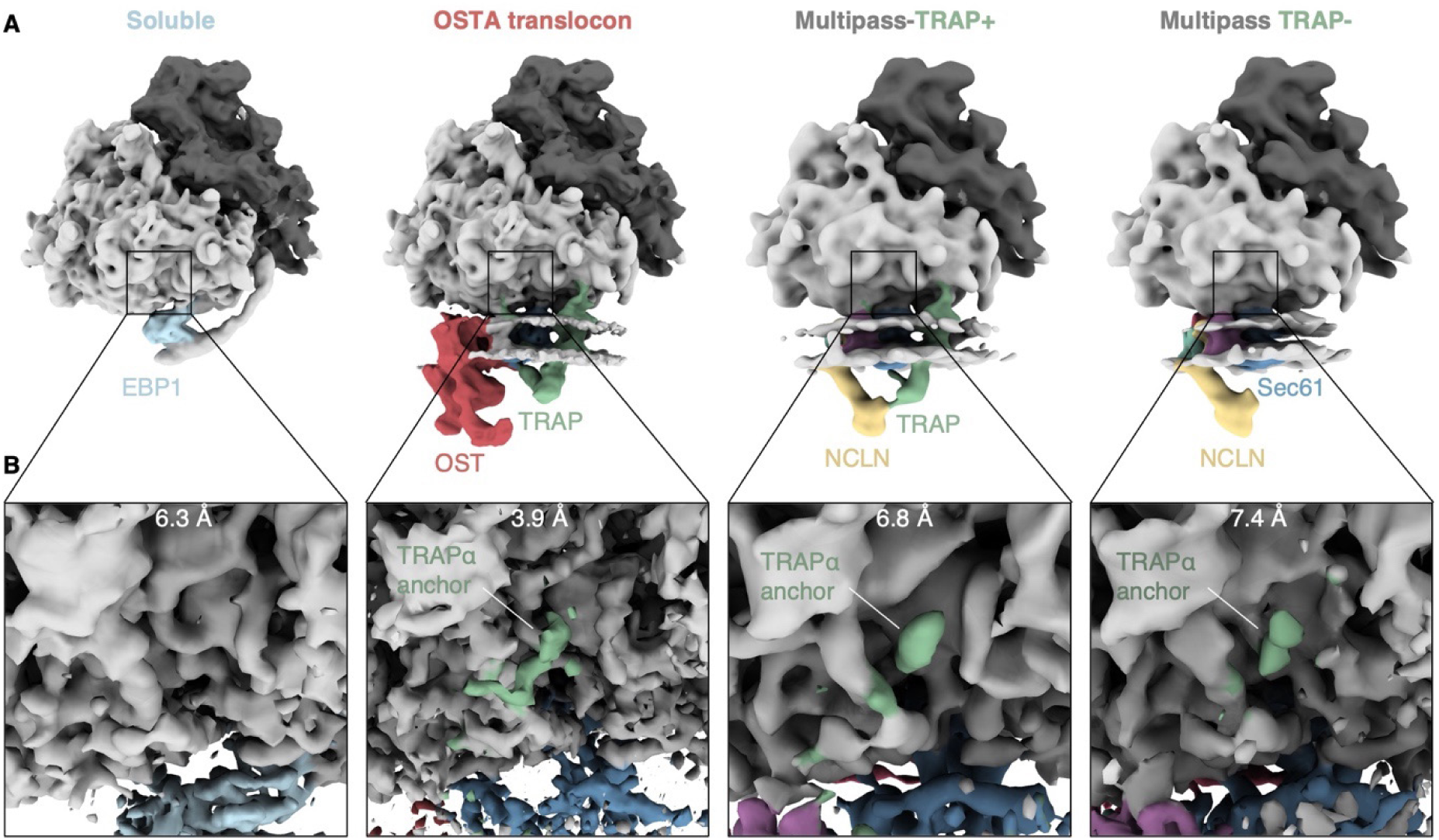
Association of the TRAPα anchor domain with distinct ribosome-bound translocon populations. (A) Back side of the ribosome associated with different soluble or ER translocon variants. Reconstructions of soluble and OSTA translocon-bound ribosomes are filtered to 12 Å and multipass-bound ribosomes to 20 Å. (B) Close-up views of the TRAPα anchor binding site. Reconstructions are filtered to their overall resolution as indicated.

Collectively, our results indicate a negative correlation of TRAP and PAT in the multipass translocon, which depends on the activity of the complex. Given that PAT correlates with actively translocating ribosomes, it is plausible that its recruitment is dependent on the presence of nascent multispanning membrane proteins. Biochemical analysis of multipass translocon assemblies engaged with defined insertion intermediates are consistent with this idea and illustrate the substrate-dependent recruitment behavior of PAT during translocation ^20^. Earlier studies demonstrated that the PAT complex subunit asterix, an intramembrane chaperone, specifically engages TMHs with exposed hydrophilic residues and releases its substrates upon correct folding ^19^. Thus, we speculate that PAT associates with the multipass-translocon-ribosome only in the presence of client TMHs with exposed hydrophilic residues.

On the opposite side of the Sec61 complex, TRAP associates preferentially with inactive multipass translocon-bound ribosomes, while its abundance in active complexes is strongly reduced. In the context of other translocon variants (Sec61-TRAP, Sec61-TRAP-OSTA), TRAP was previously considered a strictly stoichiometric accessory factor of the ER translocon ^7^. TRAP’s substoichiometric presence in multipass translocons suggests that release of TRAP may be required for efficient multipass-TMH insertion. While TRAP is known to support insertion of SPs with below-average hydrophobicity and above-average glycine-plus-proline content and contributes to safeguarding membrane protein topogenesis ^9,35^, the effect of TRAP on multipass membrane protein biogenesis has not been investigated to date. One possible scenario is that disassociation prevents TRAP-assisted insertion of TMHs into the Sec61 lateral gate.

### TRAP interacts with the BOS complex

The mechanism underlying TRAP release remains unclear since TRAP, PAT, GEL, and BOS can coexist in the multipass translocon. However, it has been suggested that the luminal segment of the BOS complex may displace TRAP at the pore exit of Sec61 ^20^. To examine the structural organization of the two accessory factors in the native membrane, we obtained multipass translocon structures containing TRAP (TRAP+/PAT– and TRAP+/PAT+, 8,477 particles) and lacking the TRAP complex (TRAP-/ PAT– and TRAP-/PAT+, 6,194 particles). In the absence of TRAP, the luminal segment of BOS adopts a tilted conformation with respect to the membrane (Fig. 4). This conformation is consistent with the orientation observed in the cryo-EM structure of the isolated TMCO1 translocon (6W6L ^19^), which also lacks the TRAP complex. In the presence of TRAP, BOS undergoes a ∼20° rotation and projects almost orthogonally from the membrane, in line with a cryo-EM structure of the multipass translocon, in which TRAP remained bound to the multipass translocon (7TUT ^20^) (Fig. 4, S1). Intriguingly, BOS would clash with the luminal domain of TRAPα when adopting the tilted conformation. Thus, conformational switching of BOS could trigger release of TRAP from the multipass translocon, as suggested earlier^20^. Vice versa, however, TRAP dissociation and recruitment may affect the conformation of the BOS complex. Further experiments will be required to determine the molecular mechanism of TRAP release and recruitment and its interplay with the BOS complex.

**Fig. 4:**
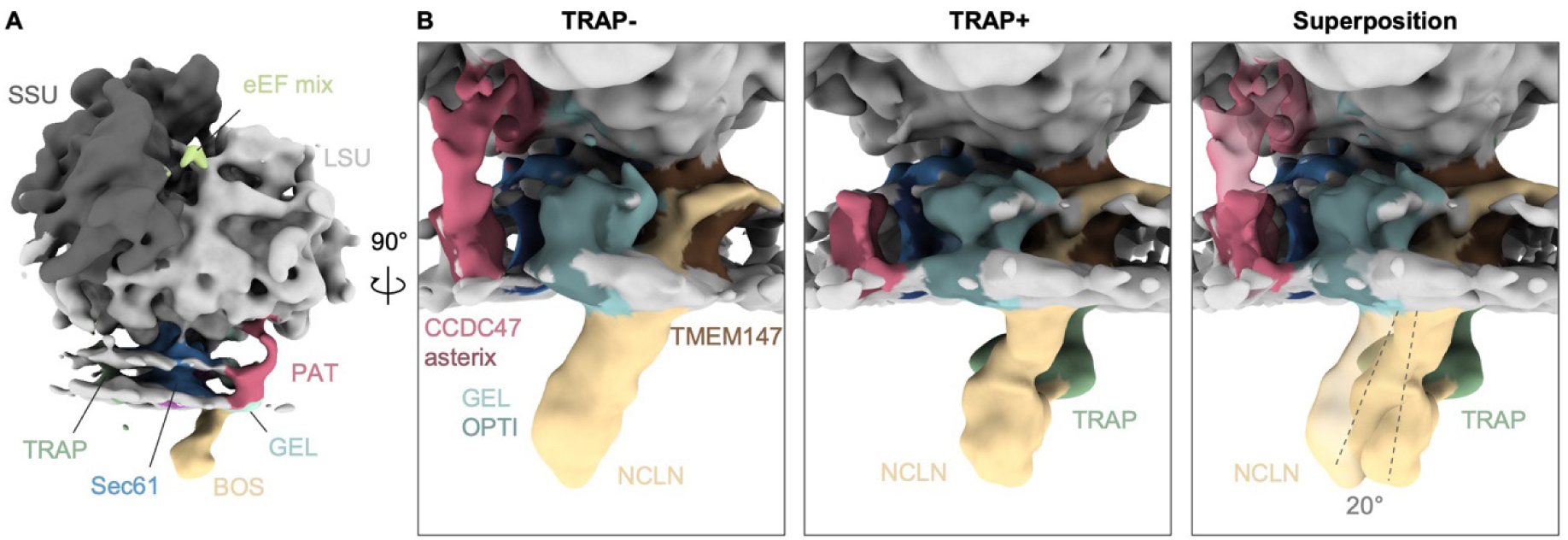
Orientation of the BOS complex is TRAP-dependent. (A) Overall structure of the ribosome-bound multipass-translocon filtered to 20 Å. (B) Close-up side views of the TRAP-lacking (TRAP-) and TRAP-containing (TRAP+) multipass translocon, and their superposition with TRAP-in transparent. The longitudinal axis of BOS is indicated as dashed line.

Subtomogram recentering and local refinement focused on the luminal densities of TRAP and BOS yielded notably improved densities with a resolution of approximately 15 Å (Fig. 5A). While the atomic models of NCLN (AF-Q969V3) and TRAP (8B6L) could be fitted into the recentered reconstruction, two small densities, each with an approximate molecular weight of 10 kDa, remain unexplained. Both densities are bound to the luminal domain of NCLN, while the membrane-proximal density interacts with the luminal domains of TRAPβ and TRAPδ (Fig. 5A).

**Fig. 5:**
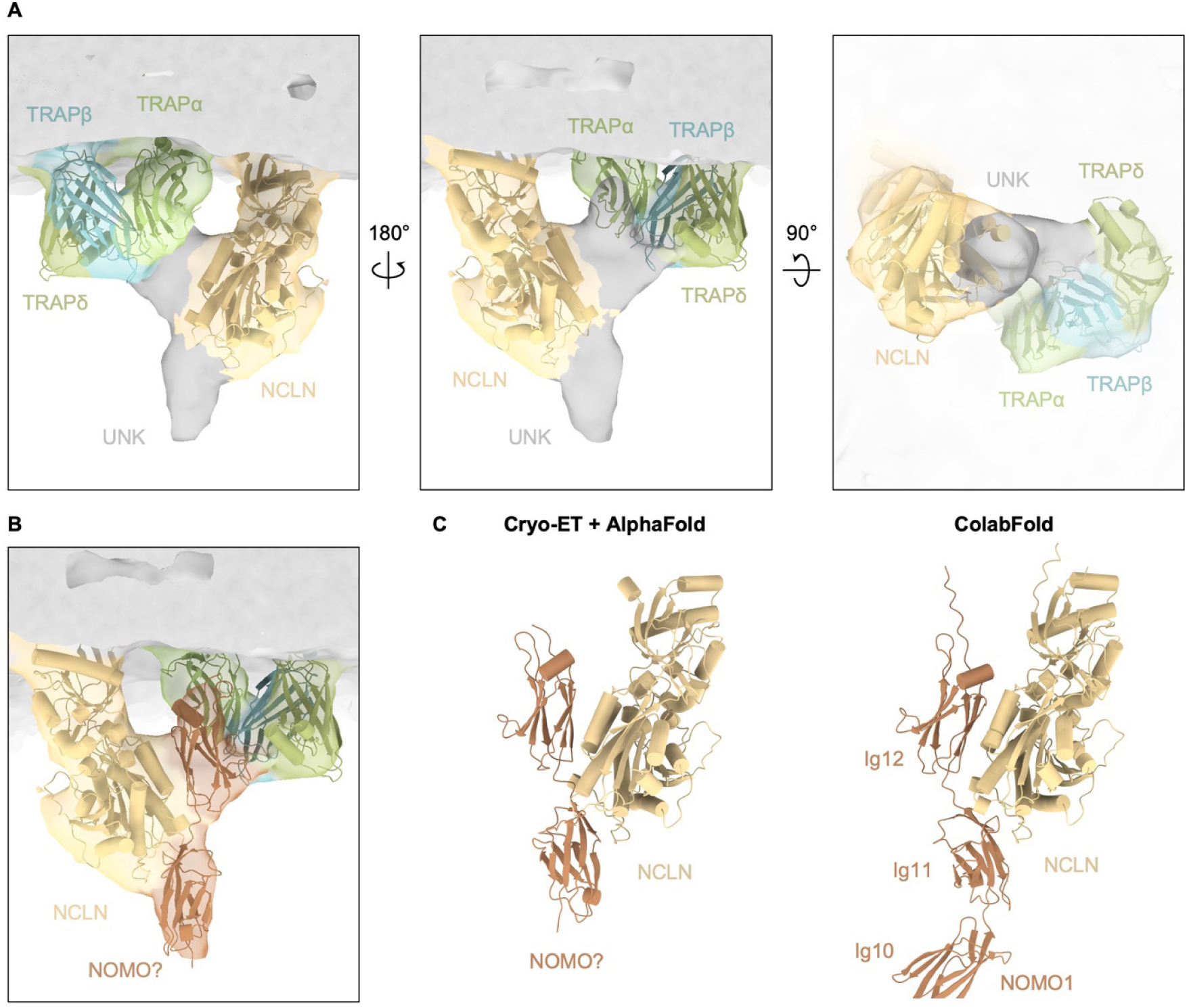
TRAP interacts with the BOS complex. (A) Recentered and locally refined reconstruction of the TRAP-containing multipass translocon. Luminal domains of TRAP (8B6L, green) and NCLN (AF-Q969V3, yellow) fit well into the density map. Two unidentified densities (UNK, grey) have a molecular weight of 10 kDa each. (B) Molecular model of the C-terminal Ig-like domain 11-12 of NOMO1 (AF-Q15155) placed into the density map of the unidentified densities. (C) Comparison of our model based on the reconstruction from (B) and the ColabFold prediction model of the NCLN-NOMO1 complex. NOMO1 domains 1-9 were removed for clarity.

An obvious candidate for the unidentified densities is NOMO, a subunit of the BOS complex ^25–27^, because was shown to be highly abundant in isolates ^19^. Among the three NOMO paralogs, NOMO1 and NOMO3 are enriched in the isolated ribosome-associated multipass translocon as detected by mass spectrometry. NOMO is a 134 kDa ER-resident single-pass membrane protein with a 124 kDa luminal domain. NOMO and NCLN were shown to associate independently of TMEM147 and the domains that mediate their interaction reside in the ER lumen ^26^. Despite its abundance in the isolate, NOMO was not modelled into the SPA reconstruction, presumably because the resolution of ER luminal densities was affected by flexibility^19^. Negative stain electron microscopy and sequence analysis of NOMO suggest an elongated shape, comprising 12 immunoglobulin (Ig)-like domains that are arranged like ‘beads on a string’ ^36,37^. AlphaFold models of NOMO paralogs are consistent with this notion ^37,38^. The two most C-terminal NOMO domains, which reside adjacent to the ER membrane-anchoring TMH, are prime candidates to explain the unknown densities and indeed fit well into the low-resolution cryo-ET reconstruction without major clashes to NCLN or TRAP (Fig. 5B). They contact TRAP at the hydrophobic cradle formed by the luminal domains of TRAPα, TRAPβ and TRAPδ, which was hypothesized to interact with translocating substrates ^34^. Thus, NOMO may occupy this site preferably at inactive multipass translocon complexes. The remaining NOMO Ig-like domains cannot be explained in the subtomogram average, which is likely due to their flexibility.

To test the hypothesis that NOMOs C-terminal Ig domains constitute the unknown density, we built a model of the NCLN-NOMO complex using Colabfold ^39^. Intriguingly, this model positions the C-terminal NOMO1 domains almost identical to our fit into the cryo-ET structure with high confidence (PAE<10, pLDDT>85%) (Fig. 5C, S2). Although experimental validation of our assembly model remains, our results strongly suggests that the luminal domain of NOMO mediates association of TRAP and NCLN in the ER lumen.

A direct interaction between TRAP and NOMO has not been reported to date. In humans and *C. elegans*, NCLN, NOMO, and TMEM147 (or their homologs) have been shown to regulate levels, subcellular localization, and subunit composition of different homo– and heterooligomeric multispanning membrane proteins ^28–30,40^, pointing to a role in membrane protein biogenesis. Moreover, NOMO has recently been shown to affect ER sheet morphology. The precise functions of NOMO, NCLN, and TMEM147 and their interplay with TRAP or other translocon accessory factors, however, remain poorly understood.

## Conclusion

Taken together, the extensive and intricate interaction network formed by Sec61-embracing accessory factors in the multipass translocon constitutes a dynamic frame allowing intermolecular communication (Fig.6). We suggest that it may regulate various processes during translocation of multispanning membrane proteins. While our results provide a comprehensive picture of the multipass translocon in its native membrane environment, further detailed functional and structural characterization in context of defined substrates is needed to understand the interplay of these specialized translocon components.

**Fig. 6:**
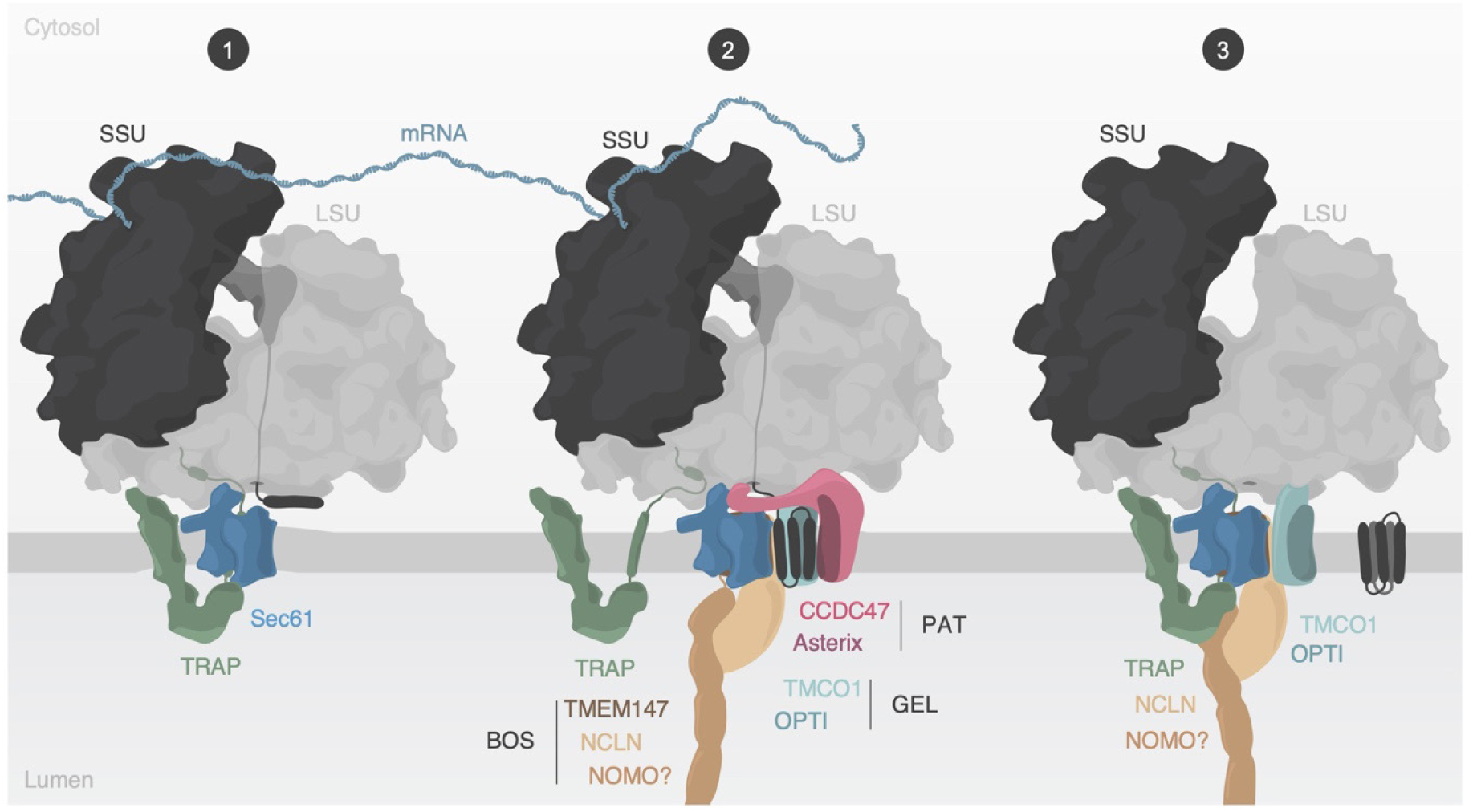
Model of multipass membrane protein biogenesis. (1) Synthesis of multispanning membrane proteins is facilitated by the ER translocon-bound ribosome. The TRAP complex engages with the translocon during the early stage. (2) Transmembrane helices of the nascent substrate are inserted into the cavity behind Sec61, while Sec61’s lateral gate and pore remain closed. The PAT complex engages with exposed hydrophilic areas of substrate proteins and the TRAP complex partially dissociates from the translocon likely to avoid TRAP-assisted insertion into Sec61’s lateral gate. The TRAPα anchor domain may remain bound to the ribosome to retain TRAP in proximity to the translocon. (3) After substrate folding and release, PAT dissociates from the ribosome and translocon, while TRAP is recruited again and interacts with the BOS complex in the ER lumen.

## Materials and methods

### Sample preparation and cryo-ET data processing

Cell culture, sample and grid preparation, data collection and processing, tomogram reconstruction, and particle localization were described previously ^31^.

### Subtomogram analysis

Subtomogram analysis of all ribosomes was described previously. In brief, the localized particles were extracted and aligned in RELION (3.1.1) ^41^. The aligned particles were further refined in 2-3 rounds in M (1.0.9) ^42^ focused on the LSU. Afterwards, newly extracted subtomograms were subjected to 3D classification in RELION to discard remaining false positives, poorly aligned particles, and lone LSUs. The remaining 134,350 particles were used for subsequent focused classification steps.

### Classification of ER translocon variants

134,350 ribosome particles were subjected to 3D classification (without reference, with soft mask, T=4, classes=20) in RELION focused on the ribosomal tunnel exit. Particles were sorted into Sec61-TRAP-bound, Sec61-TRAP-OST-bound, Sec61-multipass-bound, EBP1-bound ribosomes (soluble) and a combined class of ribosomes with ambiguous densities. Ribosomes with ambiguous densities were subjected to two further classification rounds and sorted the respective class from above until no further separation could be achieved.

We reextracted subtomograms of the multipass-translocon (14,671 particles) at a voxel size of 6.9 Å recentered onto the center of Sec61. Subsequently, subtomograms were subjected to classification focused on the ER luminal area (with reference of the entire multipass-translocon population, with soft mask, T=3, classes=2) or focused on the cytosolic domain of the PAT complex and ribosomal tunnel exit (with reference, with mask, T=3, classes=2). The TRAP-containing multipass-translocon was refined using local angular searches in RELION. Subtomograms were centered onto the ribosome again, reextracted, and averaged.

### Model fitting and prediction

TRAPα, TRAPβ, TRAPδ (8B6L), NCLN (AF-Q969V3) and luminal Ig-like domains 11-12 of NOMO1 (AF-Q15155) were fitted into the density map of the recentered reconstruction of the Sec61-TRAP-multipass translocon. The AlphaFold Colab ^43^ model prediction was build based on sequences of human NCLN (Q969V3) and NOMO1 (Q15155). The sequence of the signal peptide was removed from NCLN and NOMO prior to prediction.

### Statistical analysis

Statistical analysis was described previously ^31^.

## Acknowledgements

This work was supported by the European Research Council under the European Union’s Horizon2020 Program (ERC Consolidator Grant Agreement 724425 – BENDER) and the Nederlandse Organisatie voor Wetenschappelijke Onderzoek (Vici 724.016.001, research program TA 741.018.201, and National Roadmap for Large-Scale Research Infrastructure NEMI 184.034.014).

## Author contributions

M.G and F.F. conceived the project. M.G. performed microsome sample preparation, cryo-ET data acquisition and image analysis. M.L.C performed statistical analysis. M.G. and F.F. analysed the data and wrote the manuscript.

## Data availability

We made use of a previously published atomic models from the PDB (8B6L, 6W6L, 7TUT) and the AlphaFold Protein Structure Database (Q969V3, Q15155).

## Supplementary Figures

**Supplementary Fig. 1:**
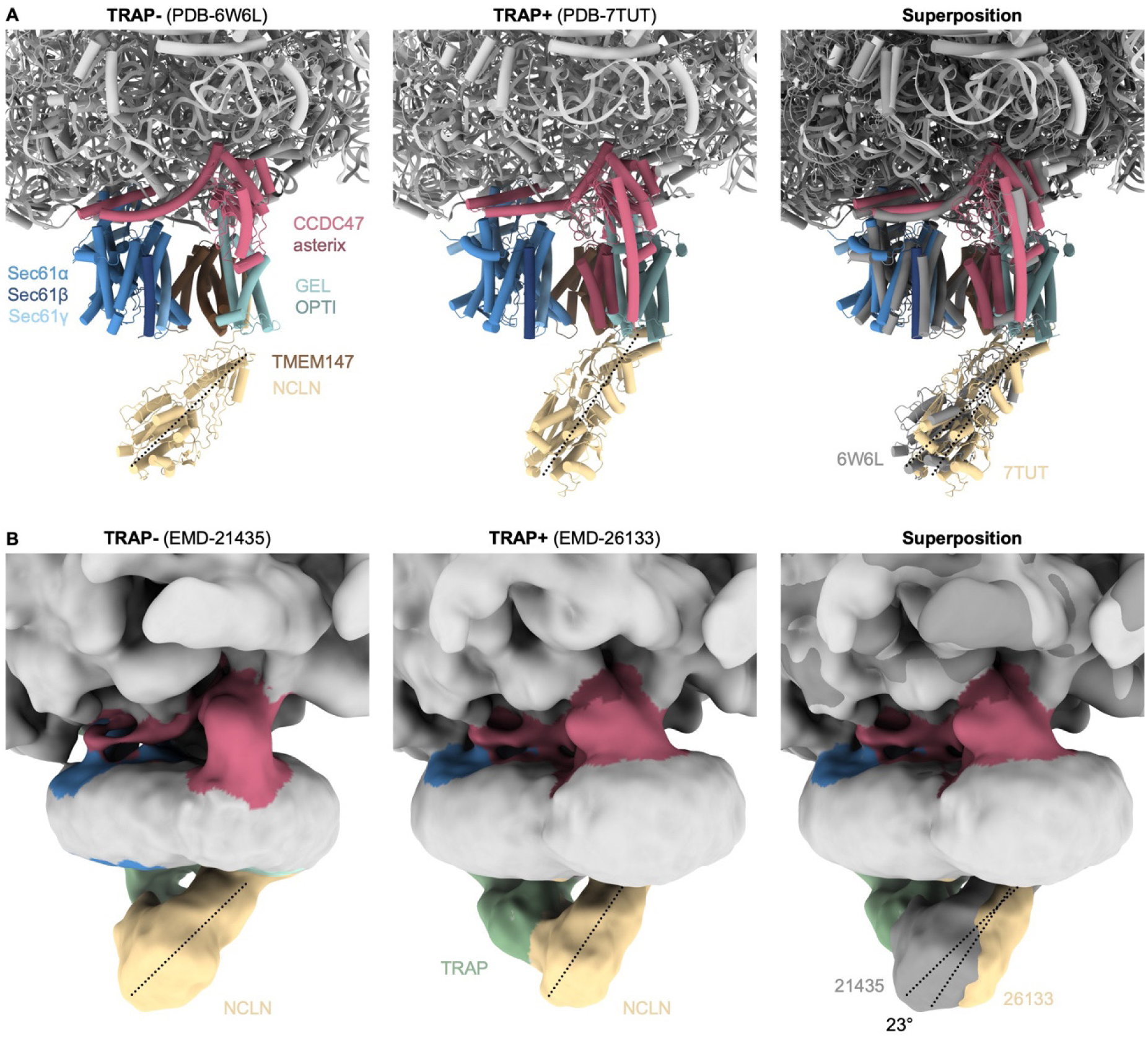
Orientation of the BOS complex in different structures. (A) Close-up side views of the TRAP-lacking (TRAP-, 6W6L) and TRAP-containing (TRAP+, PDB-7TUT) multipass translocon, and their superposition with TRAP-in dark grey. (B) Cryo-EM reconstructions corresponding to the structures from (A). Note that although a fragmented density of TRAP is present in the micelle of the TRAP-reconstruction (EMD-21435), it is disconnected from Sec61 and BOS. The longitudinal axis of BOS is indicated as dashed line. The indicated rotation angle was determined in ChimeraX.

**Supplementary Fig. 2:**
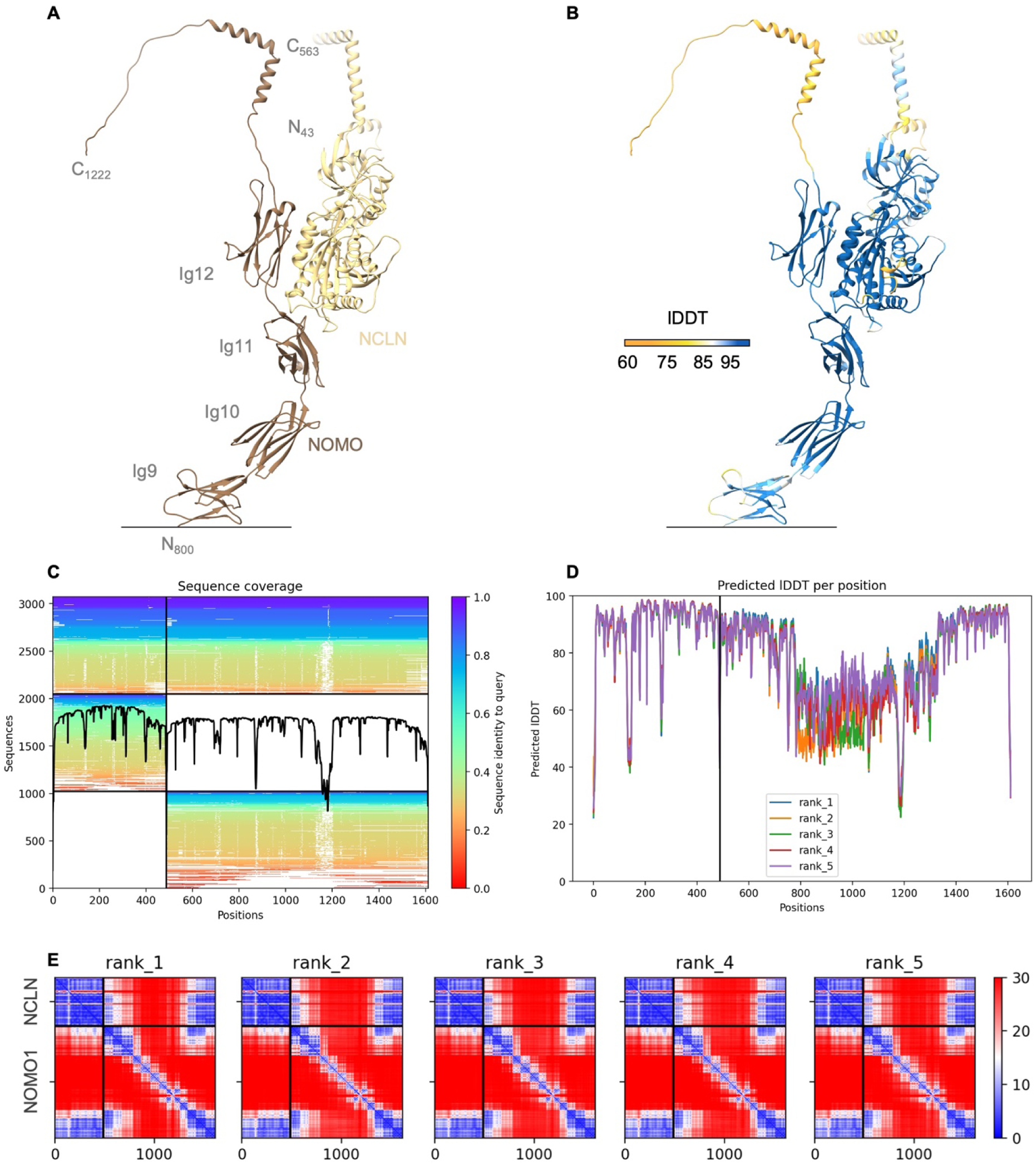
Colabfold model prediction of the NCLN-NOMO1 complex. (A) Colabfold prediction model of NCLN (full-length) in complex with NOMO1 (aa800-aa1222; Ig9-C-terminus) color-coded according to chain as indicated. Ig-like domains 1-8 were removed for clarity. N- and C-termini, corresponding residue numbering, and NOMO1 domain numbering is indicated. Signal peptides were removed prior to prediction. (B) Prediction model as in (A) color-coded according to predicted local distance difference test (pLDDT) score. (C) Sequence coverage obtained by sequence alignments generated by MMseqs2. (D) pLDDT scores per position of five model predictions. (E) Predicted aligned error (PAE) of five model predictions of the NCLN-NOMO1 complex.

**Supplementary Fig. 3:**
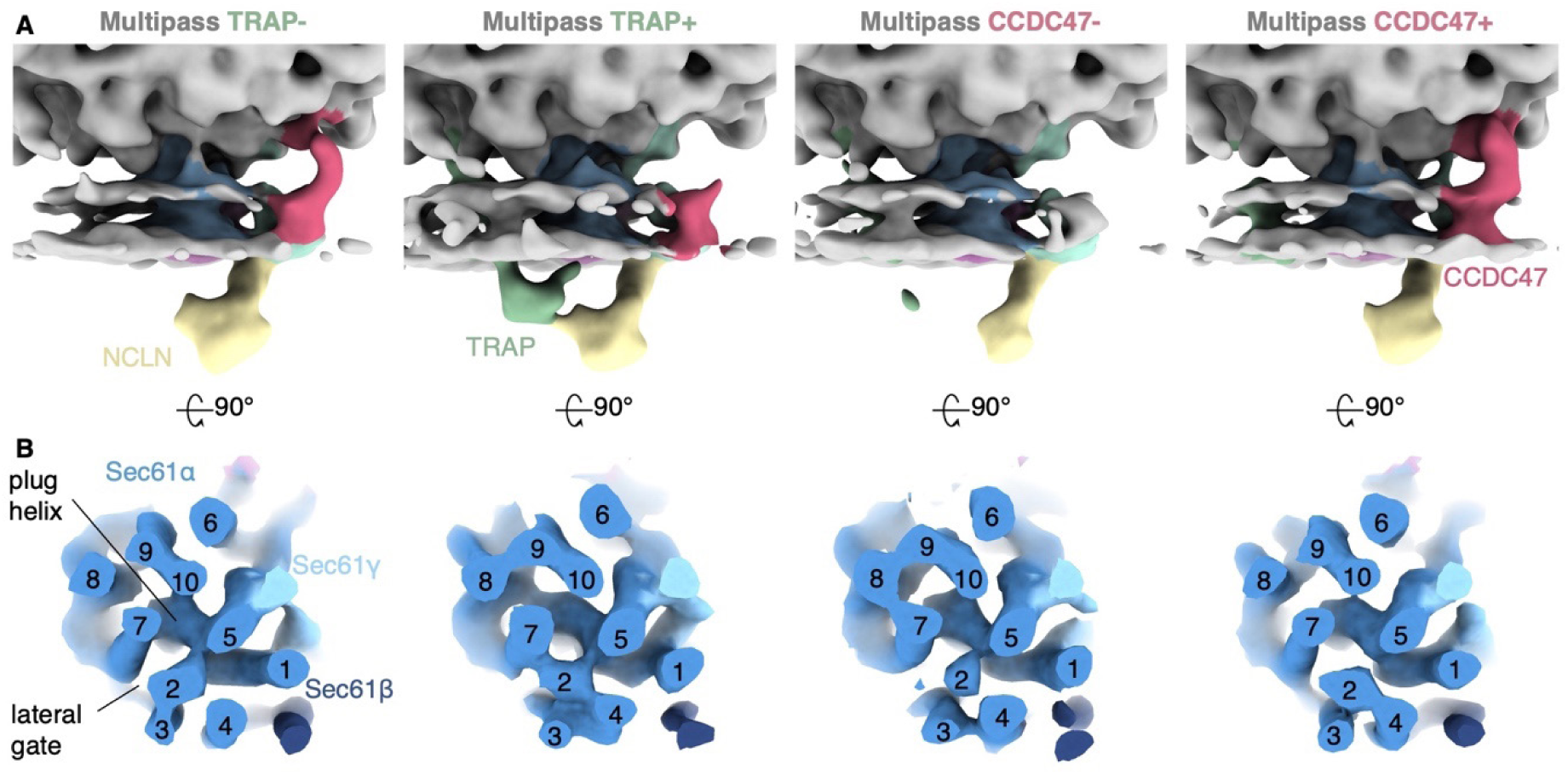
Sec61 conformational states of different multipass translocon populations. (A) Reconstructions of TRAP-lacking (TRAP-) and –containing (TRAP+), and CCDC47-lacking (CCDC47-) and –containing (CCDC47+) multipass translocon population filtered to 20 Å. (B) Top view of the corresponding map of Sec61 filtered to 8 Å. The membrane resides in the paper plane. The complex was clipped in the center of the membrane plane for visual clarity. TMH numbering is indicated.

