## Supplementary Figures for "Exploring the molecular composition of the multipass translocon in its native membrane environment"

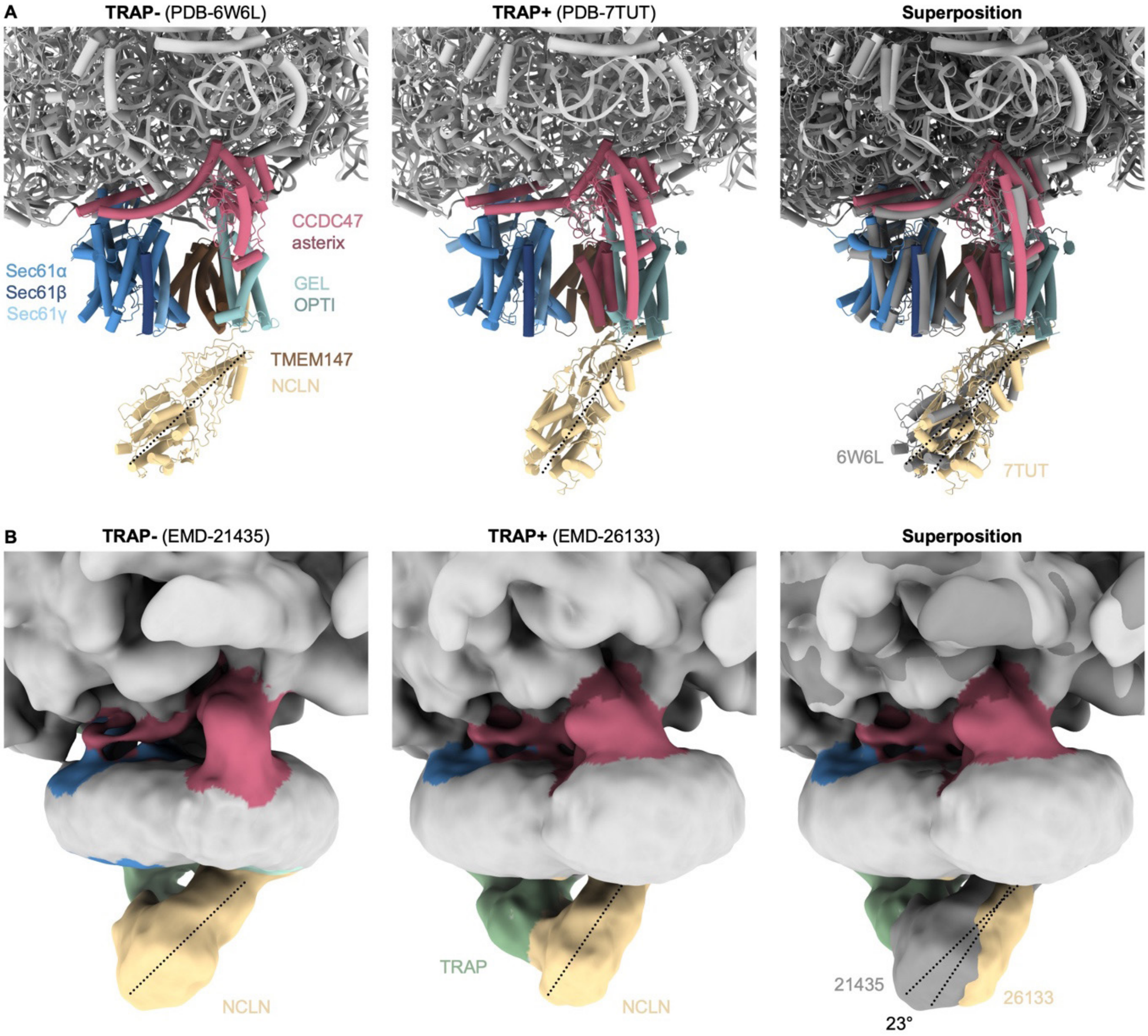


**Supplementary Fig. 1: Orientation of the BOS complex in different structures.** (A) Close-up side views of the TRAP-lacking (TRAP-, 6W6L) and TRAP-containing (TRAP+, PDB-7TUT) multipass translocon, and their superposition with TRAP- in dark grey. (B) Cryo-EM reconstructions corresponding to the structures from (A). Note that although a fragmented density of TRAP is present in the micelle of the TRAP- reconstruction (EMD-21435), it is disconnected from Sec61 and BOS. The longitudinal axis of BOS is indicated as dashed line. The indicated rotation angle was determined in ChimeraX.

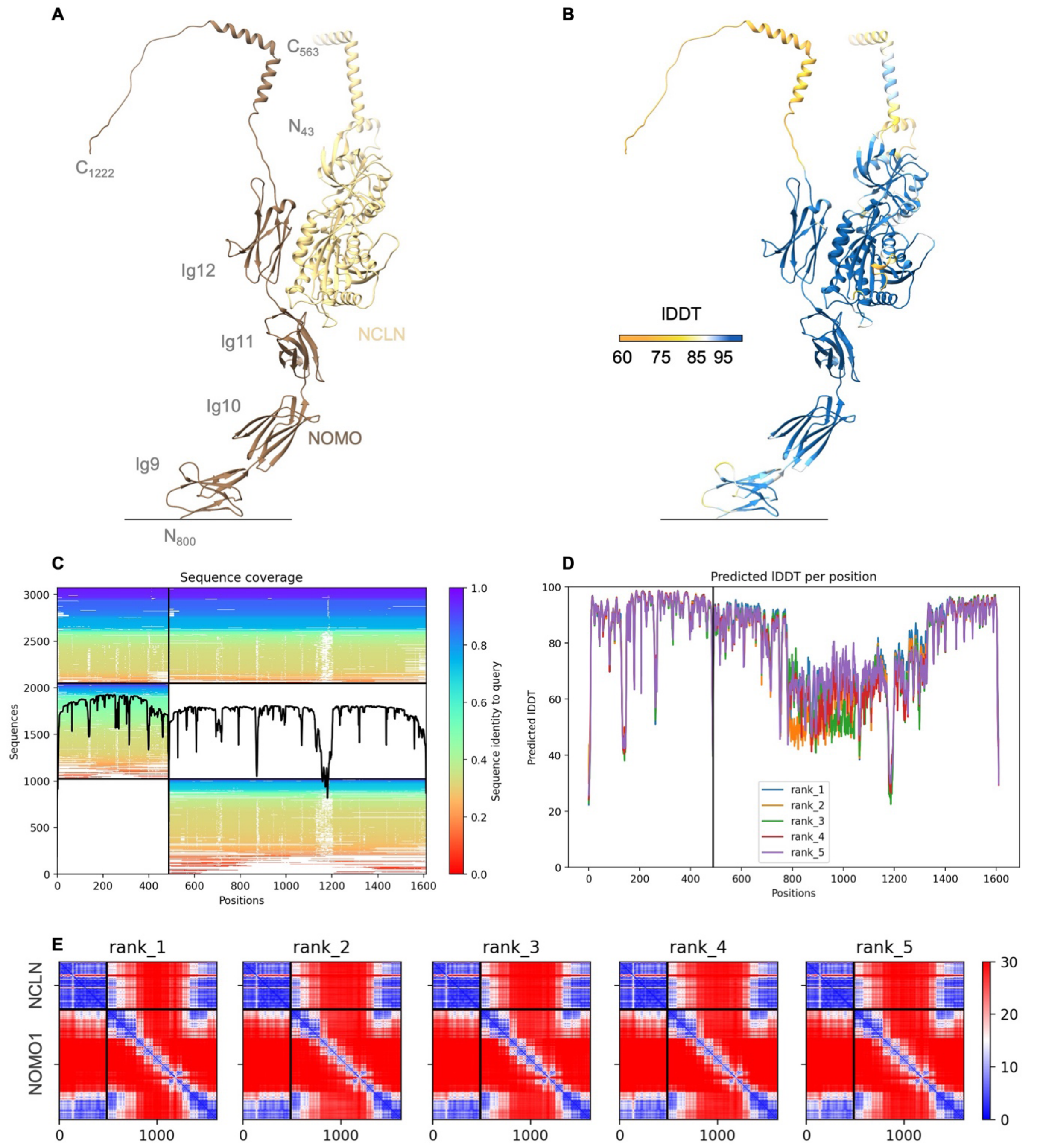


**Supplementary Fig. 2: Colabfold model prediction of the NCLN-NOMO1 complex.** (A) Colabfold prediction model of NCLN (full-length) in complex with NOMO1 (aa800-aa1222; Ig9-C-terminus) color-coded according to chain as indicated. Ig-like domains 1-8 were removed for clarity. N- and C-termini, corresponding residue numbering, and NOMO1 domain numbering is indicated. Signal peptides were removed prior to prediction. (B) Prediction model as in (A) color-coded according to predicted local distance difference test (pLDDT) score. (C) Sequence coverage obtained by sequence alignments generated by MMseqs2. (D) pLDDT scores per position of five model predictions. (E) Predicted aligned error (PAE) of five model predictions of the NCLN-NOMO1 complex.

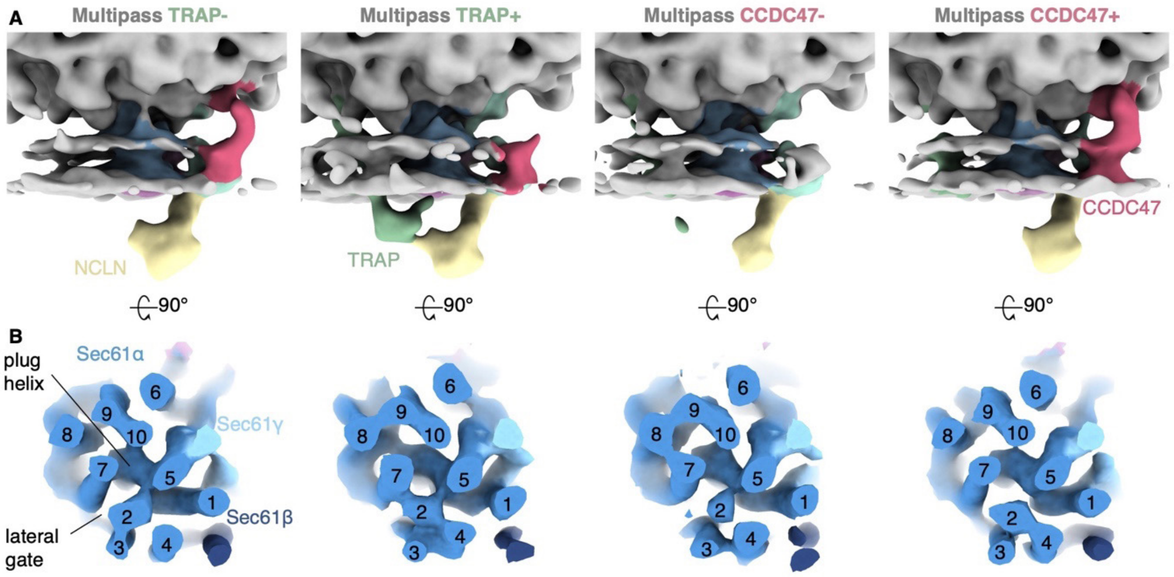


**Supplementary Fig. 3: Sec61 conformational states of different multipass translocon populations.** (A) Reconstructions of TRAP-lacking (TRAP-) and -containing (TRAP+), and CCDC47-lacking (CCDC47-) and -containing (CCDC47+) multipass translocon population filtered to 20 Å. (B) Top view of the corresponding map of Sec61 filtered to 8 Å. The membrane resides in the paper plane. The complex was clipped in the center of the membrane plane for visual clarity. TMH numbering is indicated.
